# Promoting Transparent, Fair, and Inclusive Practices in Grantmaking: Lessons from the Open and Equitable Model Funding Program

**DOI:** 10.1101/2023.11.15.567266

**Authors:** Eunice Mercado-Lara, Greg Tananbaum, Erin C. McKiernan

## Abstract

The Open Research Funders Group (ORFG) has been instrumental in promoting open and equitable scholarship to enhance academic research’s transparency and accessibility. In 2020, the ORFG formed the Equity & Open Science Working Group, leading to the launch of the Open & Equitable Model Funding Program in 2021. This program aimed to refine grantmaking processes for broader inclusivity and equitable open scholarship practices.

The program gathered a cohort of 11 funding organizations and focused on applying various proposed interventions within existing funding programs to improve grantmaking practices. The initiative highlighted the importance of targeted intervention selection and the need for delimited goal-setting, revealing challenges in implementation due to limited resources and the complexity of the processes involved.

Participants valued the pilot for its structured approach and opportunities for shared learning despite some facing complexities in fully implementing the interventions. The ORFG’s future approach involves flexible intervention selection, effective resource management, and fostering a collaborative community, aiming for a more inclusive and practical application of open scholarship principles.

## 1. Introduction

### 1.1 Why Open & Equitable Scholarship?

Open Scholarship^2^ is a movement that aims to reduce barriers to participation and incentivize collaboration in the academic research enterprise by increasing transparency, reproducibility, and accessibility of research. It encompasses various aspects such as open access, open data, and open-source software (Fecher & Friesike 2014). By making research outputs and academic discussion more widely available, open scholarship can increase the findability, accessibility, re-use, and re-distribution of research products (McKiernan 2017), thereby accelerating discovery and better addresing the big challenges of our society (Besançon, et al. 2021). In essence, open scholarship is not just about open access to scholarly outputs but also about inclusivity and the collective advancement of knowledge and the academic enterprise for a better understanding and creation of solutions to the societal challenges we face.

Nonetheless, pushing researchers toward specific sharing practices or models without considering the context of their resources can exacerbate existing inequities. Furthermore, providing adequate support for open scholarship practices is crucial; otherwise, without proper infrastructure, training, and incentives, the barriers to participation may not only persist but potentially widen the gap between well-resourced and under-resourced scholars (Ross-Hellauer, et al. 2022). Equity-related challenges can pose significant obstacles to participation^3^. This is why it is essential to tailor open scholarship initiatives to be sensitive to the diverse needs and constraints of the global research community (Chan et al. 2011).

**Table 1.** Potential Barriers to Participation in Open Scholarship, Identified from an Equity Perspective.

| Type of Barrier | Description |
| --- | --- |
| Economic | Access to open scholarly outputs often requires internet access and the use of computers or other devices, which can be expensive. Additionally, while open scholarship is often freely accessible, participating actively in it can come with costs that are prohibitive for some, especially those from low-income countries or underfunded institutions. For instance, pushing scholars towards certain open-access publishing business models, contributing data, or attending conferences can present economic challenges. |
| Educational | A certain level of guidance and training is required to engage actively with academic research. Those without this background may struggle to participate fully in open scholarship initiatives. |
| Language | Much of the scientific literature is published in English, which can be a barrier for those who do not speak the language. This can limit both the consumption of open scholarly outputs and the contribution to them. |
| Cultural | Different cultures have different approaches to sharing information, and there may be mistrust or misunderstanding of the intentions behind open scholarship. Moreover, certain cultures may restrict the sharing of knowledge that is considered traditional or sacred. |
| Technological | Not all regions have the same level of technological advancement or infrastructure. This can hinder participation in digital open scholarship platforms and limit access to online resources. |
| Legal and Policy | Intellectual property laws, data protection regulations, and institutional policies can limit the sharing of data and findings, making it difficult for researchers to participate fully in open scholarship. |
| Infrastructure | Even if researchers are willing to participate, they may work in institutions that lack the infrastructure or support systems necessary for engaging with open scholarship, such as institutional repositories or access to open-source software. |
| Bias and Discrimination | Systemic biases can affect which research is supported, whose work is published, and who gets credit for scientific contributions. This can lead to the underrepresentation of certain groups in open scholarship initiatives. |
| Resource Allocation | There can be an uneven distribution of resources within the academic community, where well-funded and established academics from prestigious institutions have more capacity to participate in open scholarship than their less well-resourced and recognized counterparts. |
| Recognition and Incentive Structures | The current academic system often rewards traditional publication and grant metrics over open scholarship, which can disincentivize researchers, especially underrepresented and early career scholars, from participating in open scholarship practices. |
| Network and Collaboration | Established researchers often have more opportunities to build networks and collaborate. Newcomers or those from marginalized groups may not have the same opportunities, which can limit their ability to engage with the open scholarship community. |

Source: Own elaboration with crowd-sourced information from the Equity in Open Science Working Group and further community engagement.

Open and equitable scholarship serves as a means to enhance the academic enterprise rather than being an end in itself. It enables a wide array of positive outcomes in the academic, philanthropic, and societal enterprise, such as inclusivity and collaboration, increased transparency, reproducibility, and public engagement, thereby strengthening scientific inquiry’s overall integrity and impact, among others observed^4^. Addressing these barriers necessitates a collaborative approach by governments, educational institutions, funders, and the broader academic community to ensure that the principles of open scholarship are genuinely inclusive and accessible.

**Table 2:**
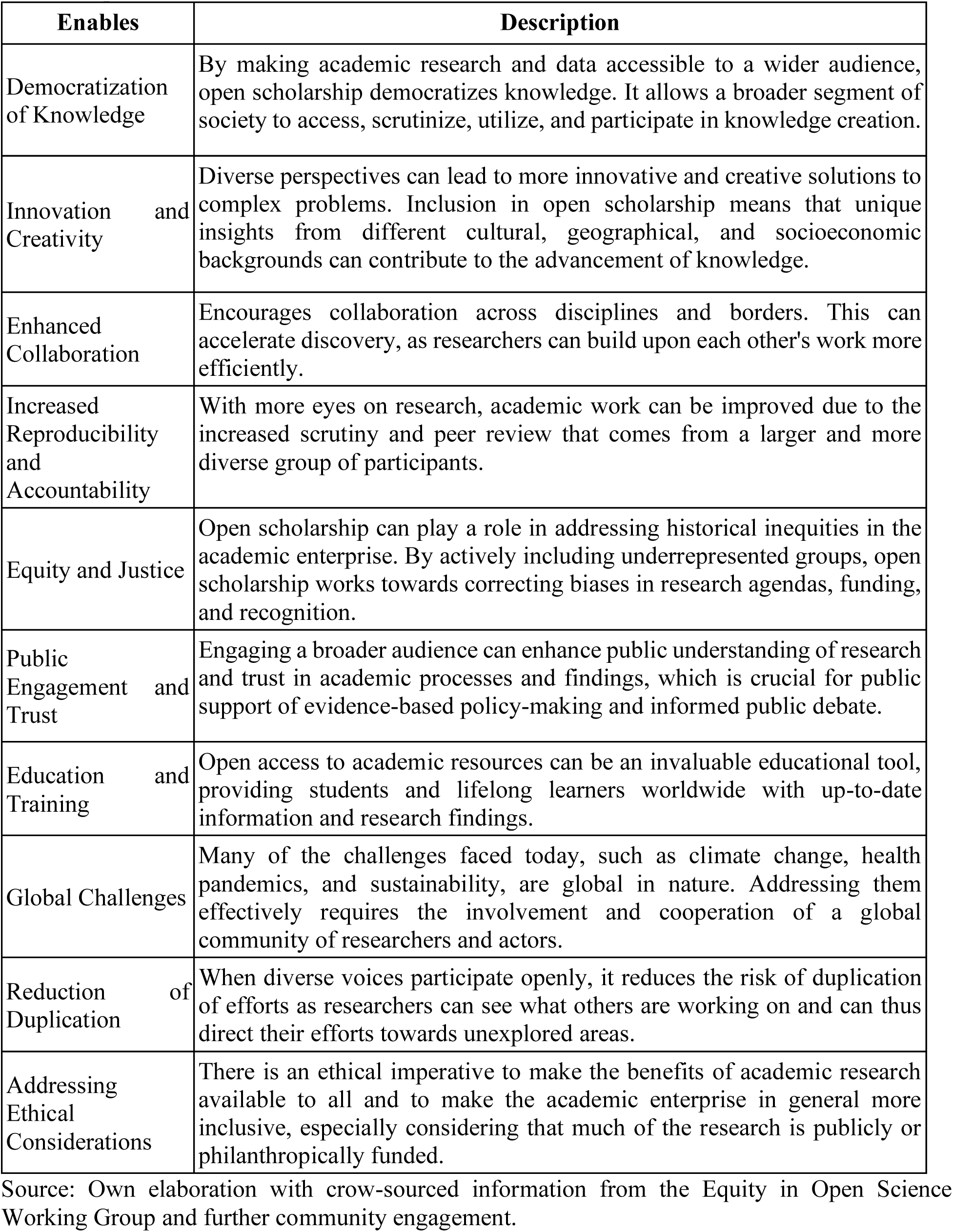
Potential Enhancements to the Academic Enterprise Achievable Through Open Scholarship.

| **Enables** | **Description** |
| --- | --- |
| Democratization of Knowledge | By making academic research and data accessible to a wider audience, open scholarship democratizes knowledge. It allows a broader segment of society to access, scrutinize, utilize, and participate in knowledge creation. |
| Innovation and Creativity | Diverse perspectives can lead to more innovative and creative solutions to complex problems. Inclusion in open scholarship means that unique insights from different cultural, geographical, and socioeconomic backgrounds can contribute to the advancement of knowledge. |
| Enhanced Collaboration | Encourages collaboration across disciplines and borders. This can accelerate discovery, as researchers can build upon each other's work more efficiently. |
| Increased Reproducibility and Accountability | With more eyes on research, academic work can be improved due to the increased scrutiny and peer review that comes from a larger and more diverse group of participants. |
| Equity and Justice | Open scholarship can play a role in addressing historical inequities in the academic enterprise. By actively including underrepresented groups, open scholarship works towards correcting biases in research agendas, funding, and recognition. |
| Public Engagement and Trust | Engaging a broader audience can enhance public understanding of research and trust in academic processes and findings, which is crucial for public support of evidence-based policy-making and informed public debate. |
| Education and Training | Open access to academic resources can be an invaluable educational tool, providing students and lifelong learners worldwide with up-to-date information and research findings. |
| Global Challenges | Many of the challenges faced today, such as climate change, health pandemics, and sustainability, are global in nature. Addressing them effectively requires the involvement and cooperation of a global community of researchers and actors. |
| Reduction of Duplication | When diverse voices participate openly, it reduces the risk of duplication of efforts as researchers can see what others are working on and can thus direct their efforts towards unexplored areas. |
| Addressing Ethical Considerations | There is an ethical imperative to make the benefits of academic research available to all and to make the academic enterprise in general more inclusive, especially considering that much of the research is publicly or philanthropically funded. |

Source: Own elaboration with crowd-sourced information from the Equity in Open Science Working Group and further community engagement.

**Table 3.** Interventions within the Open & Equitable Model Funding Program.

| Number | Intervention | Life Cycle |
| --- | --- | --- |
| 1 | Clearly articulate and communicate the links between and among open scholarship, equity, and inclusion with the organization's mission, and goals of the specific funding program to all actors within the program | Program Development |
| 2 | Develop funding programs centering on DEI (Diversity, Equity, and Inclusion) considerations |  |
| 3 | Seek legal guidance on how to maximize openness and inclusivity while ensuring appropriate protections within funding programs |  |
| 4 | Disseminate funding opportunities through an array of diverse channels | Publicity & Dissemination |
| 5 | Coordinate with diverse organizations to circulate the call to their networks and, when appropriate, create new partnerships and collaborations to co-host events geared toward specific populations |  |
| 6 | Communicate future funding opportunities to non-funded finalists |  |
| 7 | Diversify mechanisms to assist prospective applicants to decrease disparities |  |
| 8 | Simplify the application process to ensure it is not inordinately time-consuming | Application Mechanics |
| 9 | Design grant application windows and requirements to accommodate applicants with competing priorities and diverse support structures |  |
| 10 | Receive applications, and process them in languages other than English |  |
| 11 | Have a publicly available review rubric | Application Review |
| 12 | Ensure a diverse pool of reviewers |  |
| 13 | Develop a robust code of conduct for reviewers and create efficient mechanisms for reporting biased conduct during and after the review process |  |
| 14 | Train grant reviewers and program staff on how to identify and combat implicit bias |  |
| 15 | Provide concrete and actionable feedback to all applicants as part of the review process |  |
| 16 | Compensate external reviewers and other supporting personnel for their time and expertise |  |
| 17 | Consider flexible payment schedules to suit the needs of specific awardees | Strategies During the Award |
| 18 | Create supplementary grants for underresourced applicants to support services access to information, educational support for writing/curating information, and networking opportunities |  |
| 19 | Provide guidance and support to ensure grantees can disseminate their work as openly as possible |  |
| 20 | Simplify reporting practices |  |
| 21 | Define expectations of open outputs sharing in funded projects | Evaluation Metrics & Outputs |
| 22 | Endorse a comprehensive authorship and recognition mechanism |  |
| 23 | Endorse diverse scholarly products and metrics to evaluate and measure the success of funded projects |  |
| 24 | Collect demographic data from applicants and awardees for statistical purposes |  |
| 25 | Publicly share information on awarded projects |  |
| 26 | Evaluate the diversity of the grantee pool so it accurately represents the community itself |  |
| 27 | Evaluate and publicly share the impact and lessons of any the interventions implemented |  |
| 28 | Make publicly available the interventions implemented, and invite the community to provide feedback for improvements over time |  |
| 29 | Endorse track and compliance mechanisms for the program's output sharing policies |  |
| 30 | Create and adequately resource alumni networks | Alumni & Network |
| 31 | Diversify your grantees elevated in public and internal events and activities |  |
| 32 | Invite alumni as reviewers, advisory board members, and in other roles to further develop and amplify the program |  |

Source: Own elaboration with information gathered during the operation of the Open & Equitable Model Funding Program

### 1.2 Why a Model Funding Program?

Research funding organizations play a pivotal role in advancing knowledge, as they define mission objectives and allocate budgets to funding programs with distinct mandates. They can considerably influence academic incentives, fostering a research enterprise prioritizing inclusivity and Open & Equitable Scholarship practices. Some of them acknowledge the existing inequities in funding distribution, disparities in funding rates, and the tendency towards closed circles of applicant pools, which have been recognized as significant barriers to an equitable research landscape (Lauer & Roychowdhury 2021). Many funding bodies have thus engaged in equity-related initiatives as a corrective measure to help democratize access to research opportunities and reduce systemic biases (Wellcome’s Media Office 2022). This commitment to equity is an acknowledgment of funders’ role in shaping a more inclusive and diverse academic community.

Since its establishment in 2016, the Open Research Funders Group (ORFG) has cultivated a collaborative network of philanthropies to help foster research sharing policies and strategies policies, with an overall aim to enhance the accessibility, transparency, reproducibility, and reusability of scholarly outputs, including papers, data, and various other forms of research. Throughout numerous dialogues, members of the ORFG, as well as the wider funder network with which the ORFG engages, have acknowledged the need to integrate equity into the core of their open scholarship missions. They recognize that equity and open scholarship are mutually reinforcing elements; one cannot effectively function without the other.

## 2. Program Development & Characteristics

As an initial step, the ORFG joined forces with the Health Research Alliance (HRA) to establish the Equity & Open Science Working Group in 2020 (ORFG 2021a). This team, consisting of ORFG members and other scholars, scientists, open scholarship community leaders, Diversity Equity and Inclusion experts, and community builders, set out to reimagine open research, aiming for greater equity, especially for underrepresented communities. Through thorough discussions, the Working Group determined that while endorsing and bolstering open scholarship practices within their current funding structures was essential, it was only part of the solution. They recognized that funders’ grant-making capabilities were the most influential tools at their disposal to promote a more balanced and open research landscape. Therefore, the Working Group’s objective evolved to encompass not only the end products of funded studies but also the entire grantmaking process.

### 2.1 Development of the Interventions

Launched in September 2021, the Open & Equitable Model Funding Program was created to work collaboratively towards improving the way research is funded. The Working Group collaborated on an initial set of 52 possible interventions for grantmaking workflows. This group recognized that this early version was a starting point that necessitated broader input, particularly from scholars historically underserved by conventional funding mechanisms.

The ORFG proactively engaged with the academic community to refine these interventions. It hosted open community calls, which attracted around 50 participants from five countries – the U.K., U.S., Netherlands, Mexico, and Argentina. In addition to these interactive sessions, significant input was received asynchronously, allowing for a broader range of contributions from those unable to attend the live discussions. Among the contributors included scholars, funding program managers, and leaders of open projects from various sectors, such as research funding organizations, universities, and advocacy groups. The participants significantly collaborated to provide feedback, exchange experiences, and suggest ideas to improve targeted interventions, present results, and enhance mechanisms for participation. This critical exercise allowed for a deep dive into the challenges and obstacles faced, particularly from traditionally marginalized contexts, to engaging in open scholarship practices.

Through this extensive community engagement, the ORFG gathered valuable input and identified and classified a comprehensive list of barriers to engaging open scholarship practices from an equity perspective^5^. Identifying these barriers was crucial as it was fundamental in refining the interventions, so they address the specific challenges identified through community feedback and collaboration.

The ORFG also analyzed and logically structured these interventions to align them with the grantmaking life cycle stages for operational clarity and easy implementation. These stages included program development, program dissemination, application mechanics, application review, strategies during the award, evaluation metrics & outputs, and the engagement of program alumni and networks^6^.

To facilitate the practical application of these insights, the ORFG crafted detailed intervention primers, complete with a suite of actions, resources, and other materials. These comprehensive ‘how-to’ guides were designed to adapt these recommendations to various contexts. The ORFG also supplied relevant templates and tools to aid participants in implementing the strategies outlined in the primers. These enhanced interventions and accompanying guides were published in draft form on the program’s website in December 2021 (ORFG 2021b), inviting further public input.

The ORFG also created a list of potential enhancements that could be achieved by implementing open scholarship practices in academic environments from input gathered from the community. This list was instrumental in communicating the program vision to potential funders while recruiting cohort participants^7^.

### 2.2 Assembly of the Funder Cohort

After publicly posting the interventions, the ORFG began to recruit members from both the ORFG and the HRA to engage in a pilot program designed to turn them into practice. The ORFG sought program officers or executive-level staff willing to select and apply an appropriate subset of these interventions to at least one of their funding programs for at least one funding cycle, with an expectation to share their insights and experiences with the broader funder cohort. A varied group of 11 funding organizations committed to participate in this initiative (ORFG 2021c).

## 3. Pilot’s Implementation & Analysis

### 3.1 Profile of the Funder Cohort

The composition of the cohort was heterogeneous in terms of organizational size, funding capacity, and reach. Specifically, 30% of the organizations were classified as small, with a workforce under 20 employees; more than 45% were considered medium-sized, with a staff count between 21 and 100; and the remaining 25% were large organizations, employing over 100 individuals. Financially, the cohort’s annual grants ranged widely, from $5 million USD to as much as $560 million USD, and their endowments spanned from $200 million USD to $12 billion USD. Geographically, their funding efforts were distributed, with 18% focusing on regional initiatives, over 45% on national projects within the United States, and more than 27% extending their support to international endeavors. Additionally, the composition of the funding group reflected support for a broad array of research disciplines but a concentration in the biomedical research field. The representation of disciplines within the cohort was distributed as follows: 8% dedicated to the humanities, 16% to the field of education, 25% to mathematics and physical sciences, and 50% to biomedical research.

### 3.2 Profile of Participants

As a prerequisite of the pilot, it was essential for each participating organization to involve at least one senior-level individual who oversaw a funding program. This requirement highlighted the need for figures capable of offering their perspectives and steering the application of the recommendations within their specific funding initiatives. Organizations had autonomy in selecting the rest of their pilot team. This flexibility enabled them to assemble a group of support staff and other contributors aligned with their operational needs and the pilot’s goals. Such customization ensured that each organization could adeptly evaluate and refine the recommendations according to their distinct situations and organizational structures.

On average, each organization involved in the pilot included representation by more than three staff members. Among these participants, 16% held executive-level positions, including titles such as President, Vice President, or C-suite roles. Over 40% of the representatives were at the senior level, with roles like senior program officer or the equivalent, while more than 45% held mid-level positions, such as program associate or similar. Importantly, all participating organizations had the buy-in and approval for participation from their leadership, ensuring that support for the initiative was anchored at the highest level, regardless of the representatives’ positions within the organization. This widespread organizational support was crucial for each participating group’s robust engagement and meaningful contributions.

### 3.3 Program Operations & Logistics

The ORFG launched the cohort in April 2021, presenting cohort members with the entire set of potential interventions they could choose to adopt in their funding programs. Members were empowered to select the interventions they found most applicable and beneficial for their operations and missions, with the liberty to bypass those they considered less pertinent or not feasible to implement at this time.

The cohort’s monthly meetings were structured in two distinct parts. Initially, the first segment, spanning seven sessions, was dedicated to thoroughly examining each intervention, clustered around the seven distinct phases of the grant-making lifecycle detailed in section 3.1 above. These discussions aimed to deepen participants’ understanding of each intervention, as well as to nurture a community of practice in which members felt comfortable sharing their experiences and challenges.

Participating funders started the implementation of interventions from the beginning of the pilot, and it was until nearly eight months post-launch that they started sharing early results with their cohort peers. Participants began to present to their peers what was learned as they implemented their chosen set of interventions, providing real-world feedback and evidence gathered from their practical application. This approach fostered a collaborative environment for reflection and collective learning.

During the program, the cohort members received continuous support from the ORFG. They could contact the ORFG for further guidance and assistance in implementing their selected recommendations. The ORFG provided customized templates for each funder’s specific situation to make the implementation process more efficient. Furthermore, the ORFG acted as a liaison, connecting funders with experienced professionals to address particular queries and facilitate knowledge sharing and exchange of experiences. The ORFG focused on providing practical, concrete examples and best practices to guide cohort members.

Funders retained the flexibility to adjust their choices as needed, allowing them to withdraw certain interventions they had initially chosen or to expand their selection by incorporating more recommendations as their implementation process evolved. This adaptable framework was designed to accommodate the evolving needs and insights of the funders as they worked towards integrating the interventions into their funding programs.

### 3.4 Interventions Implemented

At the outset of the program, the ORFG surveyed participating funding organizations to determine which of the 32 recommended interventions they had already independently implemented within their programs. It was observed that, on average, participating organizations had about 8 of the 32 suggested interventions in place. This finding indicates that the organizations were, to some extent, aligned with the ORFG’s recommendations from the beginning, even before joining the program. Among the interventions most commonly implemented by the participating organizations prior to joining the program were^8^:

● Publicly share information on awarded projects
● Consider flexible payment schedules to suit the needs of specific awardees
● Compensate external reviewers and other supporting personnel for their time and expertise
● Collect demographic data from applicants and awardees for statistical purposes

After joining the pilot program and discussing the potential interventions with the ORFG and their fellow funders within the cohort, participating organizations then decided to add between 1 and 16 new interventions, with the majority of organizations selecting around 6. The most frequently newly added interventions were^9^:

● Disseminate funding opportunities through an array of diverse channels
● Train grant reviewers and program staff on how to identify and combat implicit bias
● Provide concrete and actionable feedback to all applicants as part of the review process
● Simplify reporting practices

During the program, the cohort members therefore focused on both enhancing their existing interventions by expanding their scope and depth (refer to Table 4), and initiating the implementation of their new selections (refer to Table 5). On average, each cohort member worked on refining or implementing around 14 interventions in total.

**Table 4.** Interventions Being Implemented by Participating Funders Prior to Joining the Program.

| **Intervention Number** | **Intervention** | **Total of Funders that Already Had It** |
| --- | --- | --- |
| 25 | Publicly share information on awarded projects | 8 |
| 17 | Consider flexible payment schedules to suit the needs of specific awardees | 7 |
| 16 | Compensate external reviewers and other supporting personnel for their time and expertise | 6 |
| 24 | Collect demographic data from applicants and awardees for statistical purposes | 6 |
| 1 | Clearly articulate and communicate the links between and among open scholarship, equity, and inclusion with the organization's mission, and goals of the specific funding program to all actors within the program | 5 |
| 5 | Coordinate with diverse organizations to circulate the call to their networks and, when appropriate, create new partnerships and collaborations to co-host events geared toward specific populations | 5 |
| 9 | Design grant application windows and requirements to accommodate applicants with competing priorities and diverse support structures | 5 |
| 26 | Evaluate the diversity of the grantee pool so it accurately represents the community itself | 5 |
| 2 | Develop funding programs centering on DEI (Diversity, Equity, and Inclusion) considerations | 4 |
| 4 | Disseminate funding opportunities through an array of diverse channels | 4 |
| 12 | Ensure a diverse pool of reviewers | 4 |
| 13 | Develop a robust code of conduct for reviewers and create efficient mechanisms for reporting biased conduct during and after the review process | 4 |
| 15 | Provide concrete and actionable feedback to all applicants as part of the review process | 4 |
| 30 | Create and adequately resource alumni networks | 4 |
| 8 | Simplify the application process to ensure it is not inordinately time-consuming | 3 |
| 11 | Have a publicly available review rubric | 3 |
| 23 | Endorse diverse scholarly products and metrics to evaluate and measure the success of funded projects | 3 |
| 31 | Diversify your grantees elevated in public and internal events and activities | 3 |
| 32 | Invite alumni as reviewers, advisory board members, and in other roles to further develop and amplify the program | 3 |
| 3 | Seek legal guidance on how to maximize openness and inclusivity while ensuring appropriate protections within funding programs | 2 |
| 7 | Diversify mechanisms to assist prospective applicants to decrease disparities | 2 |
| 18 | Create supplementary grants for underresourced applicants to support services access to information, educational support for writing/curating information, and networking opportunities | 2 |
| 20 | Simplify reporting practices | 2 |
| 27 | Evaluate and publicly share the impact and lessons of any the interventions implemented | 2 |
| 28 | Make publicly available the interventions implemented, and invite the community to provide feedback for improvements over time | 2 |
| 6 | Communicate future funding opportunities to non-funded finalists | 1 |
| 14 | Train grant reviewers and program staff on how to identify and combat implicit bias | 1 |
| 19 | Provide guidance and support to ensure grantees can disseminate their work as openly as possible | 1 |
| 21 | Define expectations of open outputs sharing in funded projects | 1 |
| 22 | Endorse a comprehensive authorship and recognition mechanism | 1 |
| 29 | Endorse track and compliance mechanisms for the program's output sharing policies | 1 |
| 10 | Receive applications, and process them in languages other than English | 0 |

Source: Own elaboration with information gathered during the operation of the Open & Equitable Model Funding Program

**Table 5.** Interventions Piloted by Participating Funding Programs.

| **Intervention Number** | **Intervention** | **Number of Funders Implementing** |
| --- | --- | --- |
| 4 | Disseminate funding opportunities through an array of diverse channels | 5 |
| 14 | Train grant reviewers and program staff on how to identify and combat implicit bias | 5 |
| 15 | Provide concrete and actionable feedback to all applicants as part of the review process | 5 |
| 20 | Simplify reporting practices | 5 |
| 7 | Diversify mechanisms to assist prospective applicants to decrease disparities | 4 |
| 8 | Simplify the application process to ensure it is not inordinately time-consuming | 4 |
| 26 | Evaluate the diversity of the grantee pool so it accurately represents the community itself | 4 |
| 5 | Coordinate with diverse organizations to circulate the call to their networks and, when appropriate, create new partnerships and collaborations to co-host events geared toward specific populations | 3 |
| 6 | Communicate future funding opportunities to non-funded finalists | 3 |
| 9 | Design grant application windows and requirements to accommodate applicants with competing priorities and diverse support structures | 3 |
| 12 | Ensure a diverse pool of reviewers | 3 |
| 21 | Define expectations of open outputs sharing in funded projects | 3 |
| 24 | Collect demographic data from applicants and awardees for statistical purposes | 3 |
| 28 | Make publicly available the interventions implemented, and invite the community to provide feedback for improvements over time | 3 |
| 1 | Clearly articulate and communicate the links between and among open scholarship, equity, and inclusion with the organization's mission, and goals of the specific funding program to all actors within the program | 2 |
| 2 | Develop funding programs centering on DEI (Diversity, Equity, and Inclusion) considerations | 2 |
| 11 | Have a publicly available review rubric | 2 |
| 17 | Consider flexible payment schedules to suit the needs of specific awardees | 2 |
| 25 | Publicly share information on awarded projects | 2 |
| 32 | Invite alumni as reviewers, advisory board members, and in other roles to further develop and amplify the program | 2 |
| 3 | Seek legal guidance on how to maximize openness and inclusivity while ensuring appropriate protections within funding programs | 1 |
| 13 | Develop a robust code of conduct for reviewers and create efficient mechanisms for reporting biased conduct during and after the review process | 1 |
| 16 | Compensate external reviewers and other supporting personnel for their time and expertise | 1 |
| 19 | Provide guidance and support to ensure grantees can disseminate their work as openly as possible | 1 |
| 23 | Endorse diverse scholarly products and metrics to evaluate and measure the success of funded projects | 1 |
| 27 | Evaluate and publicly share the impact and lessons of any the interventions implemented | 1 |
| 30 | Create and adequately resource alumni networks | 1 |
| 31 | Diversify your grantees elevated in public and internal events and activities | 1 |
| 10 | Receive applications, and process them in languages other than English | 0 |
| 18 | Create supplementary grants for underresourced applicants to support services access to information, educational support for writing/curating information, and networking opportunities | 0 |
| 22 | Endorse a comprehensive authorship and recognition mechanism | 0 |
| 29 | Endorse track and compliance mechanisms for the program's output sharing policies | 0 |

Source: Own elaboration with information gathered during the operation of the Open & Equitable Model Funding Program

On the opposite end of the spectrum, certain proposed interventions were not adopted by any cohort participants. This was not necessarily due to a perception of these interventions as unimportant; rather, they were often considered more challenging to implement, requiring more time and resources that the organizations may not have been able to commit to during the pilot phase. In this list, we have:

● Receive applications, and process them in languages other than English
● Create supplementary grants for underresourced applicants to support services access to information, educational support for writing/curating information, and networking opportunities
● Endorse a comprehensive authorship and recognition mechanism
● Endorse track and compliance mechanisms for the program’s output sharing policies.

## 4. Pilot’s Results

### 4.1 Challenges and Lessons Learned

At the conclusion of the pilot, the ORFG surveyed cohort members to gather insights on the lessons they learned, the main challenges encountered, and their overall experience in the program. The exit survey included a question to gauge participants’ perceptions of the pilot outcomes against their initial expectations. They were offered three response options: “exceeded expectations,” “met expectations,” and “fell short.” The feedback was divided; around half of the respondents (58%) felt that the pilot “met expectations,” while the other half (42%) believed it “fell short” of their expectations. Notably, none of the participants selected “exceeded expectations” as their response.

For those participants reporting that the pilot “fell short,” diverse reasons were referred to, such as:

● The turnover of representatives and staff,
● Limited time and insufficient resources to engage more thoroughly with the interventions.
● The complexity of the process, with an understanding that achieving the desired changes is a long journey, one that extends beyond the pilot’s timeframe and requires sustained effort and commitment.
● Concern that the interventions might overwhelm applicants with more work and requirements.

For those participants who reported that the pilot “met expectations,” several reasons were cited for this positive assessment:

● The interventions were well-defined and achievable, providing clear and feasible goals for the organizations to aim for.
● The structure of the pilot helped organizations allocate dedicated time and focus to the interventions, ensuring that they were given the appropriate level of attention.
● The pilot provided a framework that facilitated initiating these practices at the executive level, where strategic decisions are made.
● It offered an opportunity to collaborate with a community of like-minded individuals and organizations, all exploring similar questions and challenges.
● The experience reinforced certain ideas and practices that were initially identified when the program was launched in 2016, affirming the value of past efforts and strategies.

In the survey, the ORFG also inquired about any unexpected learnings the organizations encountered while implementing their chosen set of interventions. The responses were categorized, revealing that the most common unexpected insight was underestimating the time and resources required to enact these changes. Additionally, there were reports of resistance encountered both within various pockets of the organizations and from external sources, primarily external reviewers or review committees.

**Graphic 1:**
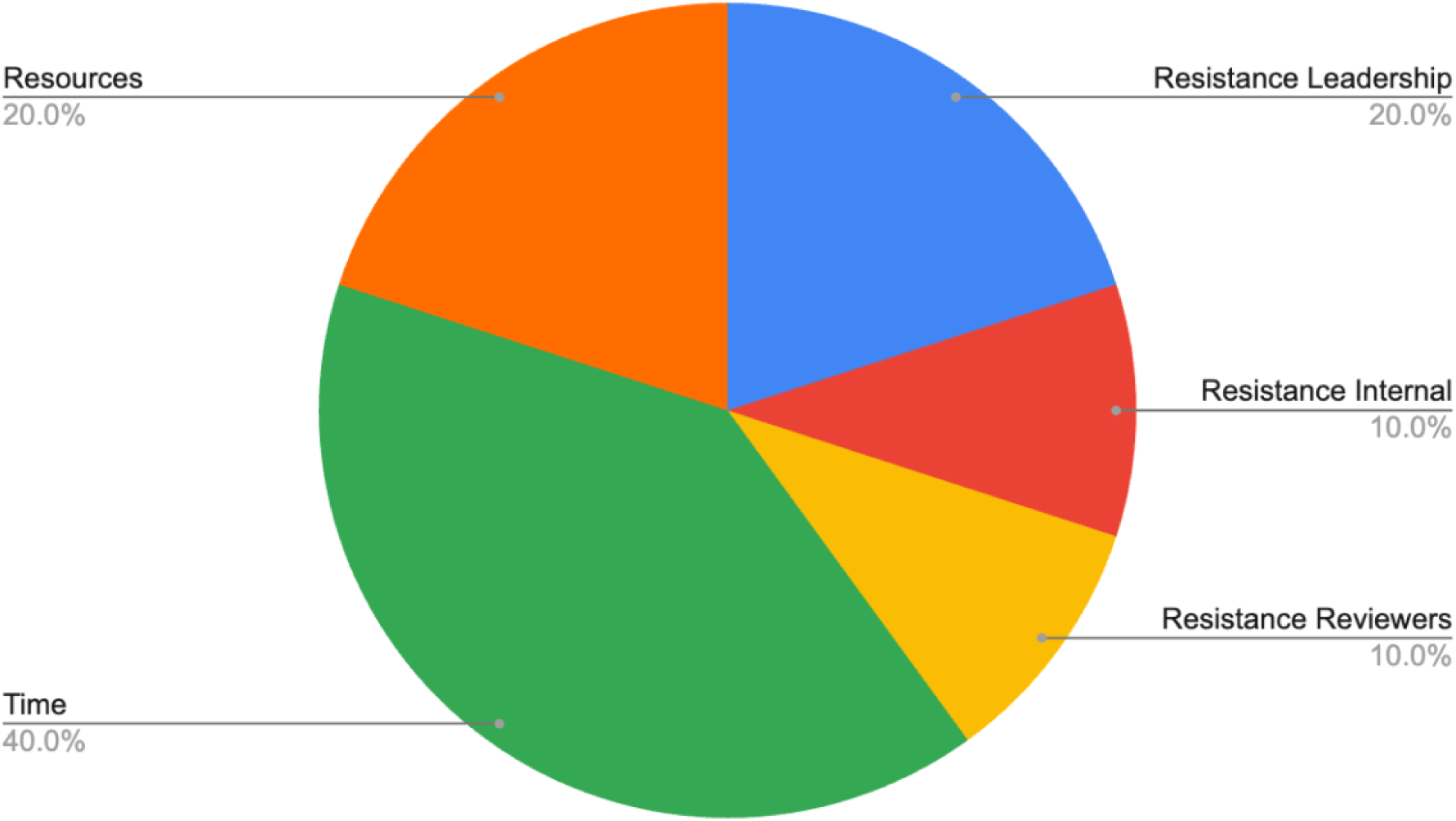
Survey Responses Percentages: What Unexpected Learnings, If Any, Did You Encounter Implementing Your Set of Interventions? Source: Own elaboration with information gathered during the operation of the Open & Equitable Model Funding Program.

Survey responses indicated a unanimous desire among the participating organizations to sustain the initiatives started during the pilot beyond its end. A majority, 58% of the cohort members, plan to extend the reach of their interventions, applying them to more funding programs within their organizations. One foundation has expressed an ambition to extend their impact further by assisting other foundations in undertaking similar work. The remaining 42% of participants intend to expand their suite of interventions, implementing a greater number within the same funding program that was involved in the pilot. This commitment reflects a robust endorsement of the pilot’s objectives and an eagerness to further integrate its recommendations into their operational frameworks.

### 4.2 Usefulness of the Pilot

Upon evaluation of their experience, 18% of the participants rated their participation in the pilot as “extremely helpful,” while a significant majority of 82% rated it as “very helpful.” This was based on a scale that included “extremely helpful,” “very helpful,” “slightly helpful,” and “not at all helpful.”

When the ORFG inquired about the most significant benefits derived from participating in the pilot, participants highlighted several key advantages, including the value of learning from the experiences of others, the resources shared, the individual guidance and personalized support provided by the ORFG team, and the benefit of committing to this work by tying it to specific deliverables and deadlines.

**Graphic 2:**
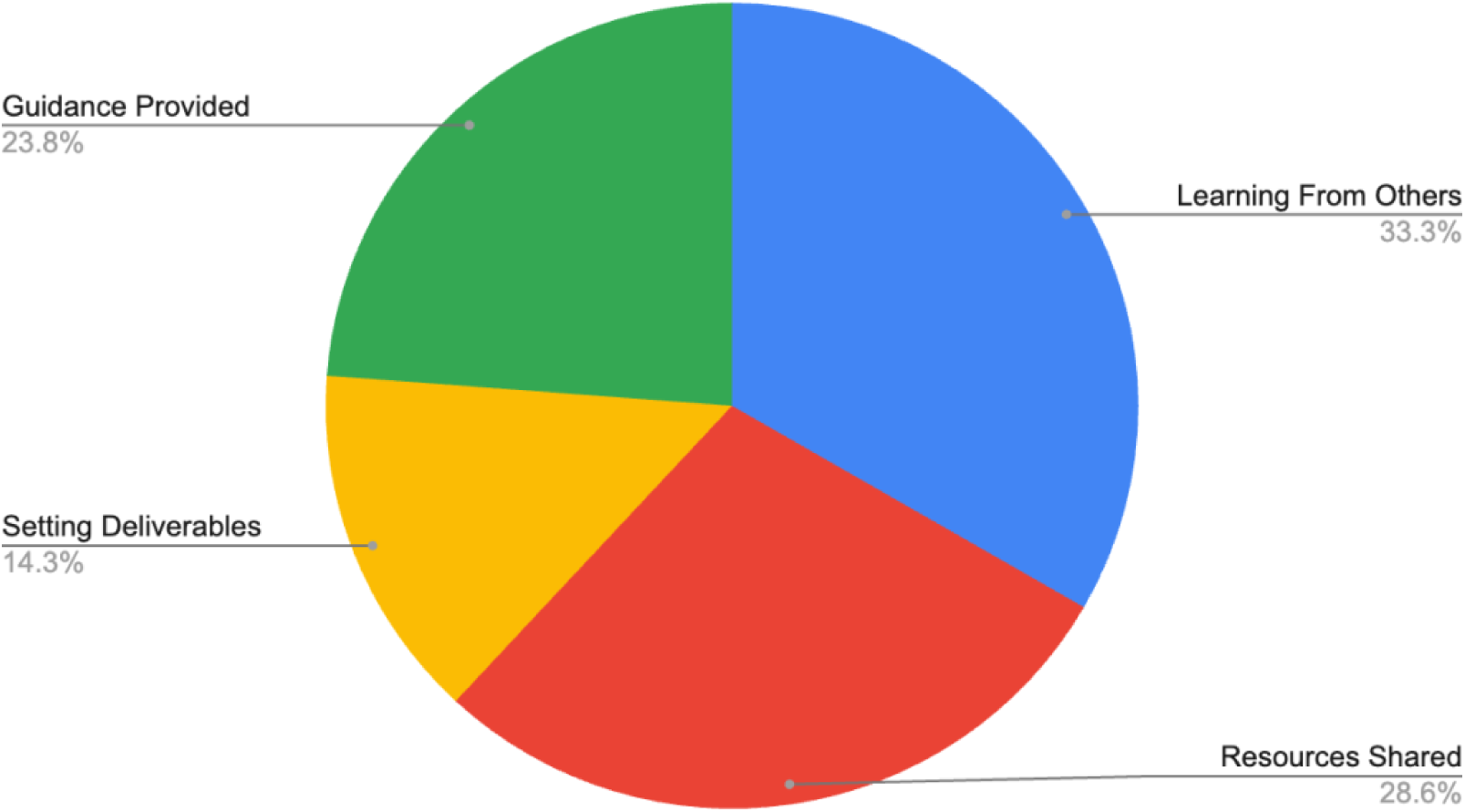
Survey Responses Percentages: What Have Been the Most Significant Benefits of Your Participation in This Pilot? Source: Own elaboration with information gathered during the operation of the Open & Equitable Model Funding Program.

### 4.3 Roadmap for Future Implementations

In light of the feedback from pilot participants, we recognize that strategic adjustments are necessary for enhancing future cohorts. These adjustments would be aimed at optimizing participant experiences and the program’s overall outcomes. They include:

1. **Expectation Management**: Manage expectations by setting realistic goals and being transparent about the potential limitations of the pilot’s impact within the given timeframe.
2. **Targeted Intervention Selection**: Encourage future participants to select and focus on a strategic and perhaps smaller number of interventions, enabling a more targeted and manageable approach to implementation.
3. **Deepen Engagement**: Introduce thematic focus into the cohort sessions to allow participants to delve deeper into specific subjects to support associated with ensuring more comprehensive discussions and clearer application to their contexts. Such as specific sessions on bias in grant review, data collection of applicants and awardees to identify diversity gaps, or development of equitable open research policies, among others.
4. **Language and Framing**: Shift the terminology from “interventions” to more accurately reflect the ongoing nature of efforts in diversity, equity, inclusion, and justice to foster a mindset of continuous cultural and structural change.
5. **Resource Management**: Address resource allocation challenges and staff turnover by providing clear guidelines on the time and resource commitments needed for each intervention, and creating contingency plans ahead of time for personnel changes.
6. **Community and Collaboration**: Strengthen the sense of community among participants through facilitated networking opportunities and shared resource platforms, encouraging peer-to-peer support and collaboration.
7. **Long-term View and Commitment**: Emphasize the long-term nature of the changes being sought, encouraging organizations to commit to sustained effort and ongoing evaluation beyond the pilot’s duration.
8. **Integration of Feedback**: Systematically integrate participant feedback into the design of future cohorts, ensuring that the program evolves to meet the needs and address the challenges identified by its participants.
9. **Diversity and Equity Commitment**: Reinforce the program’s dedication to diversity and equity in every aspect of its operation, from participant selection to the design of interventions and measurement of outcomes.
10. **Continuous Evaluation and Improvement**: Implement a cycle of continuous evaluation and improvement, using robust metrics and feedback mechanisms to inform the adaptation of the program over time.
11. **Expanding Disciplinary Participation**: Actively seek participation from funders in diverse disciplines to enrich the program with a broader range of perspectives and approaches. This expansion will enhance the diversity of insights and experiences within the cohort and contribute to developing more universally applicable and inclusive funding models.

As we chart the course for future iterations of the program, this roadmap is not just a plan but a commitment to evolve, guided by the voices and experiences of those who participate. It represents an iterative process of learning and adaptation, with each step informed by the one before it. We continue on this journey with the understanding that the path to inclusive, equitable, and open scholarship is a shared one, made richer and more rewarding by the diversity of its participants.

## 6. Acknowledgments

We recognize the valuable time and expertise contributing to this work from participating research funding organizations and the program collaborators who engaged in developing this work.

## Participating Funders

Representatives of the funding programs participating in the funders cohort (ORFG 2021c).

## Equity & Open Science Working Group Members

Tatiana Bryant. Director of Teaching, Learning, and Research Services, Barnard College.

Karen Cangialosi. Director, Every Learner Everywhere Network; Director of Open Science/Open Ed, Institute for Racially Just, Inclusive, and Open STEM.

Leslie Chan. Associate Professor in the Department of Global Development Studies and Director of the Knowledge Equity Lab at the University of Toronto Scarborough.

Elizabeth Christopherson. President and Chief Executive Officer, The Rita Allen Foundation.

Ashley Farley. Program Officer of Knowledge & Research Services, Bill & Melinda Gates Foundation.

Maryrose Franko. Executive Director, Health Research Alliance.

Monica Granados. Leadership team, PREreview.

Adam Jones. Program Officer, Science, The Gordon and Betty Moore Foundation Kari Jordan. Executive Director, The Carpentries.

Camille Thomas. Scholarly Communications Librarian, Florida State University.

Kristin Eldon Whylly. Program Officer & Senior Program Manager, Templeton World Charity Foundation.

## Partners & Contributors

Laura Ación. Project Lead / Adjunct Researcher; Ph.D.; MetaDocencia / CONICET-University of Buenos Aires, Argentina.

Robin Champieux. Director of Education, Research, and Clinical Outreach at Oregon Health & Science University Library, and co-founder of the Metrics Toolkit.

Arturo Garduño-Magaña. Open Grant Reviewers Program Manager, PREreview.

Cassandra Gould van Praag. Open Science Community Engagement Coordinator, Ph.D., Wellcome Centre for Integrative Neuroimaging, University of Oxford.

Esther Plomp. Data Steward – Faculty of Applied Sciences, TU Delft.

Daniela Saderi. Co-Founder & Director, PREreview.

Reshama Shaik. Founder & Director Data Umbrella.

Malvika Sharan. Senior Researcher, Co-lead of the Turing Way, The Alan Turing Institute. Co-Director of Open Life Science.

## Footnotes

2 We use the term “Open Scholarship” as an umbrella concept encompassing open access, open data, open educational resources, and a range of other open research and dissemination activities. Open Scholarship is used in lieu of “Open Science” to acknowledge the range of disciplines - including the arts and humanities - that engage in these practices.

3 See Table 1. Potential Barriers to Participation in Open Scholarship Identified from an Equity Perspective

4 See Table 2: Potential Enhancements to the Academic Enterprise Achievable Through Open Scholarship

5 See Table 1. Potential Barriers to Participation in Open Scholarship Identified from an Equity Perspective

6 See Table 3. Interventions Within the Open & Equitable Model Funding Program

7 See Table 2: Potential Enhancements to the Academic Enterprise Achievable Through Open Scholarship.

8 For the full list of interventions already in place within participating funding organizations, see Table 4. Interventions Being Implemented by Participating Funders Prior to Joining the Program.

9 For the full list of interventions implemented within the Model Funding Program, see Table 5. Interventions Piloted by Participating Funding Programs.

